# Increased upper-limb sensory attenuation with age

**DOI:** 10.1101/2020.09.17.301739

**Authors:** Manasa Parthasharathy, Dante Mantini, Jean-Jacques Orban de Xivry

## Abstract

The pressure of our own finger on the arm feels differently than the same pressure exerted by an external agent: the latter involves just touch, whereas the former involves a combination of touch and predictive output from the internal model of the body. This internal model predicts the movement of our own finger and hence the intensity of the sensation of the finger press is decreased. A decrease in intensity of the self-produced stimulus is called sensory attenuation. It has been reported that, due to decreased somatosensation with age and an increased reliance on the prediction of the internal model, sensory attenuation is increased in older adults.

In this study, we used a force-matching paradigm to test if sensory attenuation is also present over the arm and if aging increases sensory attenuation. We demonstrated that, while both young and older adults overestimate a self-produced force, older adults overestimate it even more showing an increased sensory attenuation. In addition, we also found that both younger and older adults self-produce higher forces when activating the homologous muscles of the upper limb.

While this is traditionally viewed as evidence for an increased reliance on internal model function in older adults because of decreased somatosensory function, somatosensation appeared unimpaired in our older participants. This begs the question of whether the decreased somatosensation is really responsible for the increased sensory attenuation observed in older people.

**New and Noteworthy:** Forces generated externally (by the environment on the participant) and internally (by the participant on her/his body) are not perceived with the same intensity. Internally-generated forces are perceived less intensely than externally generated ones. This difference in force sensation has been shown to be higher in elderly participants when the forces were applied on the fingers because of their impaired somatosensation. Here we replicated this finding for the arm but suggest that it is unlikely linked to impaired somatosensory function.

## Introduction

The position of one’s arm is monitored by sensory organs such as skin receptors or muscle proprioceptors. This information is then processed in light of a top-down organization where expectations and prior knowledge influence how the stimulus is perceived (Kok et al. 2012; de Lange et al. 2018). In essence, sensorimotor integration is a process in which the central nervous system integrates different sources of information (sensory and prior information) and transform them into motor actions (Machado et al. 2010). This processing allows humans to differentiate between internal (produced by our own movement) and external stimuli (Blakemore et al., 1998). As a result, our body perceives the sensory consequences of its own movements less intensely than the same stimulus produced by the external environment. This decrease in intensity of the perception of the self-produced stimuli is called sensory attenuation (Blakemore et al. 2000; Brown et al. 2013; Wolpert et al. 1995a) and relies on the connection between sensorimotor areas and the cerebellum (Kilteni and Henrik Ehrsson 2020).

Sensory attenuation (also termed sensory cancellation) is a widespread phenomenon that applies to different types of movements (saccades, vestibuloocular reflex, force, etc.), to perception (Cao et al. 2017; Klever et al. 2019; Niziolek et al. 2013) and is observed in many species (Sillar and Roberts 1988; Webb 2004). For instance, there is evidence of attenuation of responses to self-generated sounds in mice (Rummell et al. 2016). Flying insects need to be able to distinguish self-induced stimulation (such as rotation of the visual field caused by tracking a target) from externally imposed stimulation (such as visual rotation due to air disturbances) if they are to use the latter for flight stabilization (Dickinson and Muijres 2016; Webb 2004). Electric fish need to distinguish between perturbation of the surrounding electric field is due to a predator or due to their own movements (Kirk 1985). Sensory attenuation might also explain why humans cannot tickle themselves (Blakemore et al. 2000; Wolpert et al. 1995b), and why sounds produced by an external agent always seem louder than sounds produced by us (Klaffehn et al. 2019).

Another consequence of sensory attenuation is the tendency to underestimate the force that individuals produce in force matching tasks (Palmer et al. 2016; Shergill et al. 2003; Wolpe et al. 2016). In such tasks, participants are asked to reproduce an external force with one hand applied on the other hand (target force, e.g. 2N). They typically produce more force that they should (self-produce force, e.g. 3N) while judging that the target and the self-produced forces have the same intensity. This phenomenon is referred to as over-compensation and is a behavioral consequence of sensory attenuation.

To capture the causal relationships between our actions and their sensory consequences, the brain makes use of an internal forward model (Blakemore et al. 2000; Franklin and Wolpert 2011; Shadmehr et al. 2010; Sommer and Wurtz 2008; Wolpert et al. 1995a; Wolpert and Miall 1996). Such internal model takes a copy of the motor commands sent to the muscles (efference copy or corollary discharge) as input and outputs the predicted sensory consequences. When a sensation is internally generated (e.g. by our own movement), the internal model predicts its sensory consequences (Blakemore et al. 2000; Bubic et al. 2010; Cullen et al. 2011; Wolpert et al. 1995a) and uses this prediction to attenuate the sensory effects of the produced movement (Blakemore et al. 2000; Sato 2008; Wolpert et al. 1995). Externally generated sensations are not associated with any efference copy and are therefore perceived differently.

By attenuating the sensory consequences that are due to self-produced movement it is possible to accentuate the sensation of events caused by external agents (Moore et al. 2009). Sensory attenuation has been linked to the sense of agency, which is the perception that the observed movement has been internally generated (Kilteni and Ehrsson 2017; Moore et al. 2009). It is based on the comparison between the expected sensory consequences of the movement and the actual sensations of it (Brown et al. 2013; Moore et al. 2009; Weiss et al. 2011). If these match, the movement will be considered as internally-generated. A sense of agency over movements that generate sensation seems to be necessary for sensory attenuation; sensory attenuation does not occur if the movement and sensation are correlated, but the relationship is not perceived as causal (Brown et al. 2013; Desantis et al. 2012; Gentsch and Schütz-Bosbach 2011).

Studies have shown that sensory attenuation increases with age (Klever et al. 2019; Wolpe et al. 2016). For instance, in force matching tasks, when young and old participants experience an external force on their finger, older participants applied higher self-produced forces than younger participants. This increased overcompensation with aging might stem from age-related changes in one of the two sources of information used for sensory attenuation: sensory feedback or internal model predictions. Furthermore, the balance between these two streams of information has been shown to rely on Bayesian integration (Ernst and Banks 2002). That is, both streams are weighted in function of their relative reliability (Körding et al. 2004; Orban de Xivry et al. 2013). Given that the reliability of sensory information decreases with aging (Dunn et al. 2015; Goble et al. 2009; Ranganathan et al. 2001), it has been suggested that older adults rely more on the predictive stream (i.e. on their internal model) (Wolpe et al. 2016). In addition, some studies point to the fact that internal model function might be not be affected by aging (Heuer et al. 2011; Vandevoorde and Orban de Xivry 2019), but this is still debated (Bernard and Seidler 2014).

To our knowledge, the only two studies that evaluated age-related changes in the motor domain did so at the fingers (Klever et al. 2019; Wolpe et al. 2016). Yet, sensory attenuation has been reported for upper limbs as well (Logan et al. 2019). In this study, we want to investigate whether a larger sensory attenuation in older participants can be detected in other limbs and decided to focus on the upper limbs. We hypothesized that older adults will have a higher sensory attenuation over the arm due to increased reliance on internal models. In addition, we tested the possibility that sensory attenuation is modulated by the group of muscles sensing and producing the force. To do so, we compared the amount of overcompensation when homologous or non-homologous muscles are involved in sensing and producing the forces as network controlling homologous muscles have a particular connection as evidenced by mirroring activity during unilateral movements (Beaulé et al. 2012). We hypothesized that this connection would increase with age (Shinohara et al. 2003), hence we examined it in both young and older adults.

## Methods

Thirty-five young adults aged 18-35 years and thirty-five older adults aged 55-75 years, participated in experiment 1. Thirty-one young adults aged 18-35 years and thirty older adults aged 53-75 years participated in experiment 2. Both experiments were approved by the Ethics Committee of the University Research UZ/KU Leuven (Study number: S61179) and performed according to the guidelines of the Declaration of Helsinki. All subjects provided their written consent prior to their participation. The Edinburgh handedness questionnaire (Oldfield 1971) was used to confirm self-reported right-handedness. All participants were screened with a general health and consumption habits questionnaire. None of them reported a history of neurological disease. Older adults were assessed using the Mini-Mental State Examination (Folstein et al. 1975) for general cognitive functions. All older adults scored within normal limits (score>= 26).

### Setup

Participants were asked to grab the handles of a robotic manipulandum (KINARM End-Point Labs(tm), BKIN Technologies, Kingston, ON Canada). Their hands were hidden from view and reflected as two white cursors. These cursors were displayed on a screen placed tangentially above a mirror and were reflected by it. Because the mirror was halfway between the handle and the screen, the cursors appeared to be positioned at the same position in space as the hands. All experimental conditions were programmed in MATLAB-Simulink (Mathworks, Natick, MA, US). The force exerted on the handles were measured by built-in force transducers. Position and force data were sampled at 1000 Hz.

### Experimental paradigm

The Force Matching task implements the Method of Adjustment, in which participants adjust the level of the stimulus to match a previously presented stimulus (Wolpe et al., 2016). Two red circles appeared on the screen and participants had to reach to them and to maintain their hand position inside them. The color of the circles turned green to indicate that the hand cursors were positioned inside them. The right handle was then locked at that position in order to eliminate movements of the right handle throughout the experiment. The left circle turned then blue to indicate the start of force perception period. During the force perception period, the left hand was pushed rightwards (+X direction) by the robot with a force of 4, 6 or 8N (target force). The force was ramped up over 1s, maintained constant during 2s and then ramped down during 1 second. Participants were asked to resist the force and stay inside the circle. A safety region was included between the two hands. If participants did not resist the force enough and if, as a consequence, the left hand went above half the distance between the two targets (i.e. between the original positions of the left and right hands), the force was turned off and the trial was restarted. Three target forces of 4, 6 and 8 N were presented in pseudorandom order. Ten trials were provided for each level of force.

At the end of this phase, the right circle (above the right hand) turned blue to indicate the start of force reproduction period. In this phase, the participants had to control the robot in order to produce a force on the left hand that matched the force experienced during the force perception period. The reproduction phase differed in function of the condition.

In the slider condition, the right circle became a rectangular shape of 20 cm of height and 2 cm of length. Participants could produce force on the left hand by moving the dot located within the rectangle up (slider up) or down (slider down). The position on the slider was mapped to the force on the left handle (Fig 1A 2a. Slider). In experiment 1, subjects were given a maximum of 6 seconds to match the perceived force and were asked to apply the matched force until the end of the reproduction phase. In experiment 2, they had unlimited time but had to signal verbally to the experimenter when they had matched the target force. The experimenter ended then the reproduction phase by clicking on a button.

**Fig 1:**
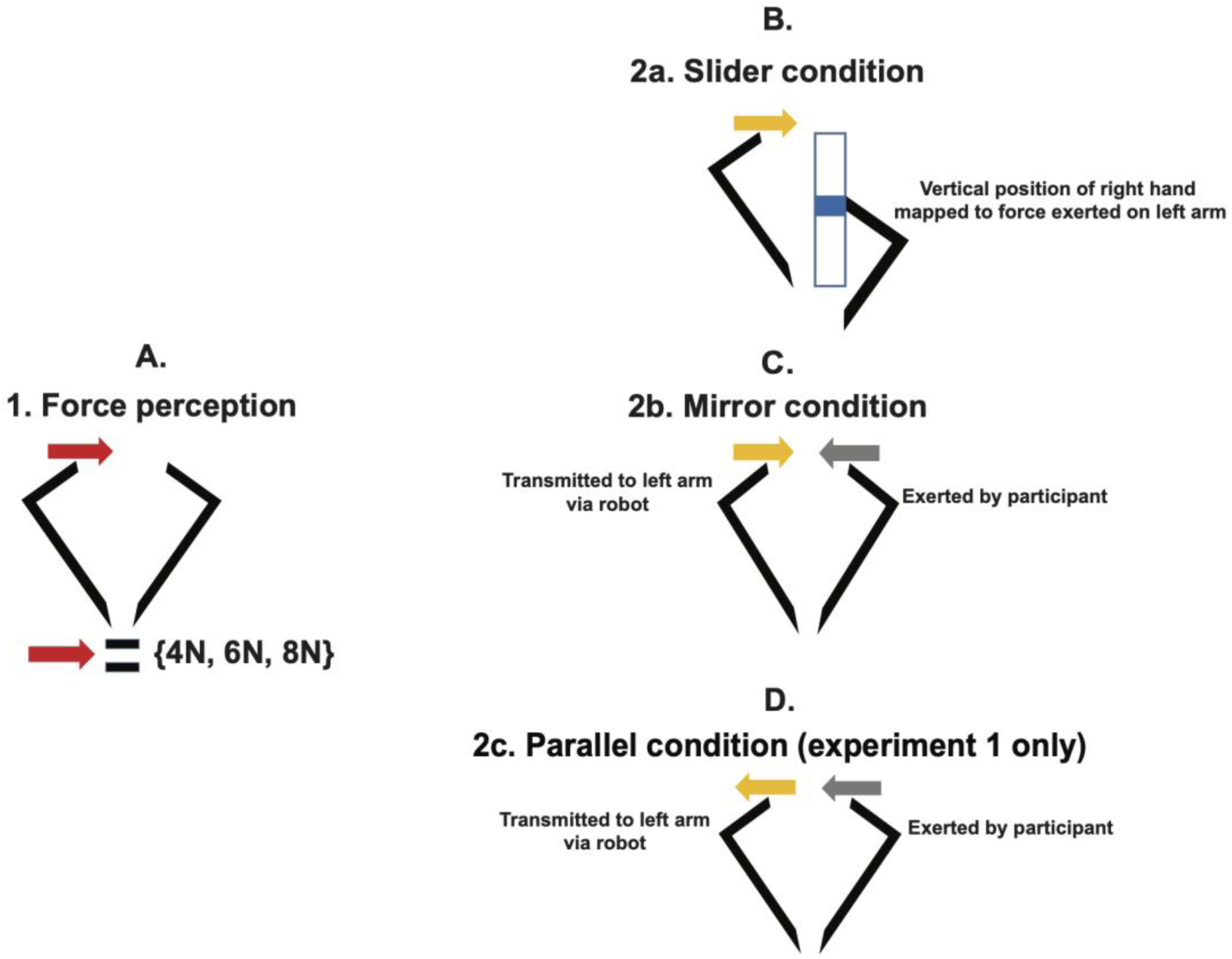
Experimental blocks of the study. Panel A: A target force that pushed the left arm to the +X direction was presented for 2 seconds (force perception phase), with a ramp-up and ramp-down of 1 second each. Participants were asked to counteract this force and judge the level. The force perception phase was followed by a second phase (force reproduction phase) where the participants were asked to reproduce the force that they perceived on their left hand. This phase differed across the three possible experimental conditions (panels B, C and D). In the slider condition (panel B), participants matched the target force by moving the right arm up or down on a slider. The position of the hand/slider was mapped to a certain level of force and transmitted to and felt on the left arm. There were also two direct Conditions: Mirror (panel C) and Parallel (panel D). In the mirror condition (panel C), participants matched the target force by applying a force to the right handle using the right arm. This was transmitted and felt on the left arm in the +X direction as shown by the arrows. In the parallel condition (panel D), participants matched the target force by applying a force to the right handle using the right arm. This was transmitted and felt on the left arm in the -X direction as shown by the arrows.

The slider condition was designed to estimate somatosensation and evaluate sensory biases in the force-matching task as the movement of the right hand was only indirectly matched to the force produced on the left hand. In contrast, there were two conditions where the force exerted by the right hand was directly mapped to the force felt in the left hand.

In the mirror condition, participants had to match the target force by exerting a force with the right hand on the right handle, in the –X direction. This produced force was transmitted to and felt on the left handle in the +X direction. In experiment 1, subjects were given a maximum of 6 seconds to match the target force and were asked to apply the matched force until the end of the reproduction phase. In experiment 2, they had unlimited time but had to signal verbally to the experimenter that they had matched the target force. The experimenter ended then the reproduction phase by clicking on a button. This condition required the activation of non-homologous muscles of the arm (biceps for the right arm to produce the force and triceps for the left arm to resist the force). The objective of this condition was to test if activation of non-homologous muscles of both arms had an effect on the perception of self-produced forces.

The direct parallel condition differed from the mirror condition in the mapping between the force produced on the right hand and the force felt in the left hand and in the instructions. In the parallel condition, the produced and felt forces were in the same direction. That is, if the right hand produced a force in the –X direction, the force produced on the left hand was also in the –X direction (Fig 1A 2c. Parallel). Furthermore, while the target force was felt in the +X direction, the participants were instructed to match the force in the –X direction. This condition was only used in experiment 1. The parallel condition requires the activation of homologous muscles of the arm.

The direct and slider conditions were counterbalanced across participants for both experiments. In experiment 1, the order of the mirror and parallel conditions was also randomized.

Before experiment 1, participants were given 9 practice trials, in each condition. In experiment 2, we programmed further instructions for each stage of the task on the screen for participants and increased the number of training blocks. First, there was a practice block where participants only felt the target forces. Next, a “play” block was provided where participants could apply the force on the right handle and feel it on the left one. The third block was a practice block that involved all stages of the task.

In both experiments, subjects were also ensured breaks in between conditions and blocks in order to prevent fatigue.

### The position matching task

To assess proprioceptive abilities, we also tested N=69 (34 young and 35 old, experiment 1) and N=56 (30 young and 26 old, experiment 2) subjects on an arm position matching task (Dukelow et al. 2010; Fuentes and Bastian 2010). Subjects were instructed to relax and let the robot move the right arm to 1 of 9 different spatial locations. When the robot stopped moving, subjects were asked to move their left hand to the mirror location in space i.e mirror-match the position of the robot. Subjects notified the examiner when they completed each trial and the examiner then triggered the next trial. Target locations were randomized within a block. Each subject completed 6 blocks for a total of 54 trials.

### Data processing

All the data collected were analyzed in MATLAB (Mathworks, Natick, MA, US). For experiment 1, produced forces were calculated as the average of the force measured by the force transducer of the left handle (which is very similar to the force produced by the participant on the right handle in direct conditions) between 1s and 3s after the start of matching phase (Fig 2). For experiment 2, the produced forces were calculated between 1s after the start matching phase and until the verbal cue of the participant when the Go button was pressed. Target forces were taken as 4, 6 and 8 N which were same as the commanded forces from the robot. Since the subject resisted the forces, the actual forces perceived during the force perception were very slightly higher or lower than the target forces of 4, 6 and 8 N.

**Fig 2.**
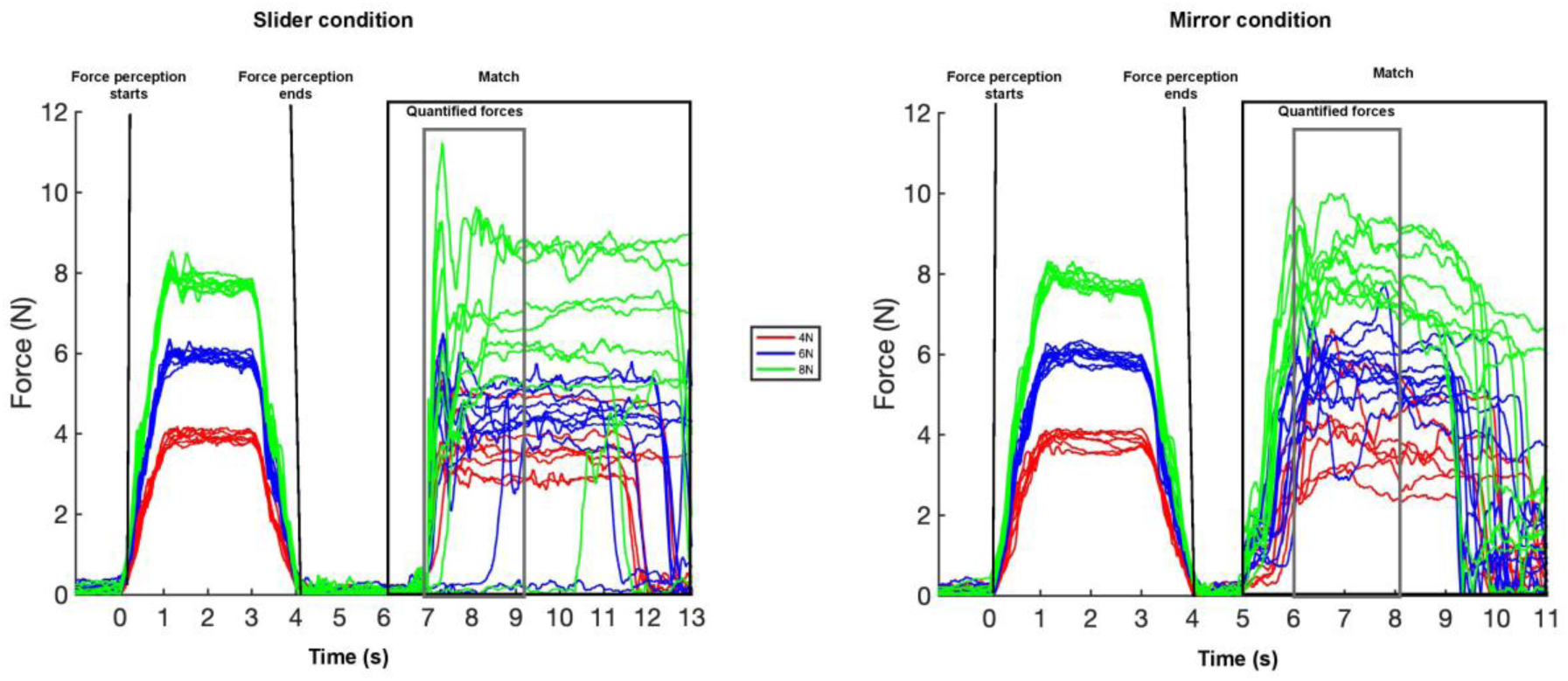
Force profile of one participant across all three levels of force and two conditions. The force profile is represented for all trials from one participant across two different conditions (left panel: slider; right panel: mirror). Each color corresponds to a different target force (4, 6 or 8 N) that was presented to the participant during the force perception phase. The X-axis represents the number of seconds since the start of the force perception period. The Y-axis shows the force measured by the force transducer of the left hand.

For each trial separately, we computed the force error as the difference between the produced force and the target force. This quantity was expressed in Newton. We then computed the normalized overcompensation as the difference in force error between the direction condition (mirror or parallel) and the slider condition. In other words, we used the slider condition as the control condition.

### Data Analysis

All values (calculated average force and force error values used in our statistical analyses) were those averaged across valid trials. Trials where the forces were not resisted enough and where the hand went into the safety region were excluded. We rejected 6.7 % of trials in the slider condition and 3% in the direction conditions for experiment 1. In experiment 2, these percentages amounted to 6.8% and 1.3%, respectively.

All analyses were performed in Matlab (Mathworks, Natick, MA, US). For slider conditions, we calculated the average between the slider up and slider down trials. In experiment 1, there was no slider down condition for one older subject. In experiment 2, there was no slider down condition for 12 older subjects.

In the paper, we report the mean (across trials) of the average force value from each trial. In the supplementary material, we also report the median (across trials) of the average force values from each trial (supplementary note 1) and the mean (across trials) of the maximum force value from each trial (supplementary note 2) for each participant, condition and experiment.

#### Analysis 1

To test for differences between the two age groups across all three force levels in experiment 1, we used a 2-way analysis of variance (ANOVA) with force errors as the dependent variable, age group as the between-subject factor and levels of forces (4, 6 and 8N) as within-subject factor. We performed this test for each condition (slider, parallel, mirror) separately.

#### Analysis 2

To test for difference in performance between age groups across all three force levels and all three conditions for experiment 1, we used a 2-way analysis of variance (ANOVA) with the three levels of forces and three conditions (slider, mirror and parallel) as within-subject factors. We performed this test separately for young and older adults

#### Analysis 3

To test for differences between the two age groups across all three force levels and all three conditions in experiment 1, we used a 3-way analysis of variance (ANOVA) with the age as the between-subject factor and levels of forces and conditions as within-subject factor, with the force level as the dependent variable.

#### Analysis 4

To test for differences between the normalized overcompensation in the two age groups across all three force levels and all three conditions in experiment 1, we used a 3-way analysis of variance (ANOVA) with the age as the between-subject factor and levels of forces and conditions (mirror vs. parallel) as within-subject factor. We also performed a t-test against zero for each condition (mirror and parallel) to test whether mean normalized overcompensation was higher or lower than zero. We performed the t-test separately in young and older adults.

#### Analysis 5

To test for differences between the two age groups across all three force levels in experiment 2, we used a 2-way analysis of variance (ANOVA) with the age group as the between-subject factor and levels of forces as within-subject factor. We performed the same test for each condition.

#### Analysis 6

To test for difference in performance between each age groups across all three force levels and all three conditions for experiment 2, we used a 2-way analysis of variance (ANOVA) with the three levels of forces and two conditions as within-subject factors. We performed this test for young and old adults separately.

#### Analysis 7

To test for differences between the two age groups across all three force levels and both conditions in experiment 2, we used a 3-way analysis of variance (ANOVA) with age group as the between-subject factor and levels of forces and conditions as within-subject factor.

#### Analysis 8

To test for differences between the normalized overcompensation in the two age groups across the mirror condition in experiment 2, we used a 1-way analysis of variance (ANOVA) with the age as the between-subject factor.

## Results

In the force-matching task, participants had to reproduce with their right arm a target force that they perceived earlier with their left arm. During the force perception period, participants experienced the target force for 2 seconds, with a force ramp up and ramp down for 1 second while trying to maintain their hand in a given position (Fig.2). During the force reproduction period, they exerted a force against the right handle of the robotic manipulandum, which was transmitted and felt on the left arm (direct condition). As shown in Fig.2 for one participant, the produced forces were generally higher than the target forces in most of the trials. In addition, this was observed across all levels of forces (Fig 2, red, blue and green lines). The level of produced force in the direct condition was compared to the control condition where the action of the right arm was indirectly linked to the force transmitted to the left arm. In the slider condition (Fig 2, left panel), participants produced the force with their right arm by moving the right handle up or down like a slider. In the direct condition (Fig 2, right panel), the produced force corresponded to the force that the participant exerted on the right handle.

Across all participants, in the slider condition, we observed that both older and young adults were able to scale the forces that they produced with the level of target force but systematically undershot the target forces across all three levels of forces. (Fig 3A and 3B). This contrasts with the observation that older participants exerted higher forces than young adults in the direct conditions (Fig 3D-E, G-H). While younger participants produced less force than the target force during the reproduction phase (Fig. 3D and 3G), the average reproduced force of older participants in the mirror and parallel conditions was higher than the target force in all but one case (8N target force in the mirror condition, Fig 3E and 3H). In addition, the produced forces appear to be larger in the parallel than in the mirror condition for both the young (Fig.3D vs. Fig.3G) and the older participants (Fig.3E and 3H).

**Fig 3.**
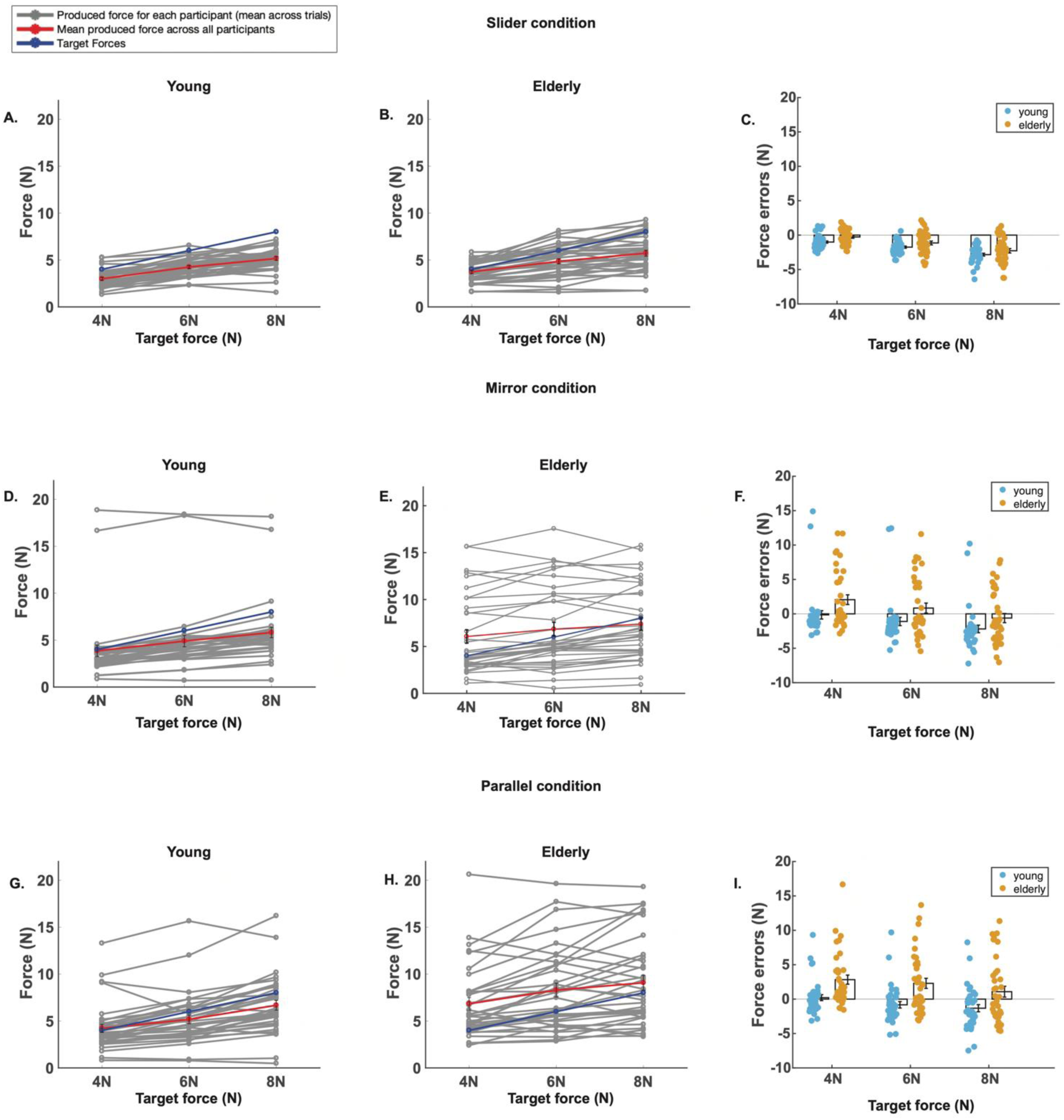
Experiment 1: comparison of exerted forces and force errors of both age groups across the three conditions. Each row corresponds to a different condition (top row: slider; middle row: mirror; bottom row: parallel). The average force exerted by the participants from the young (left column) and old group (middle column) is presented. The gray traces represented the average force for each individual participants. The red trace represents the group average. The blue trace corresponds to the target force. The X-axis shows the different force levels. Y-axis is the produced Forces (N). The third column present the average force errors (N=35/group) for both age groups and each force level. Black rectangle and error bars of 3C, 3F and 3I represent the mean and standard error respectively. Each dot is the average of all trials for each force level and for each participant.

### Force errors are lower in younger than older adults in the Slider condition

To quantify these differences between age groups, we analyzed the mean force errors (difference between produced and target force, see data processing) between young and old participants across the three levels of forces for each condition separately, starting with the slider condition.

Participants from both age groups produced a force lower than the target force in the slider condition, leading to negative force errors (Fig 3C). Force errors were closer to zero in older (Fig 3C, represented by orange dots) than young adults. That is, their undershoot was smaller than that of young adults (Analysis 1, main effect age, F(1,68)=4.8, p=0.03, *η*^2^_p_ =0.1402). In addition, participants exhibited increasing negative force errors with increasing levels of forces (Analysis 1, main effect level of force, F(2, 136)= 137.5, p<0.0001, *η*^2^_p_ =0.857). This was consistent in young and older participants, as we did not detect a between-group difference in the scaling of the force errors with increasing target force (Analysis 1, level of force x age, F(2, 136)= 0.311, p=0.73, *η*^2^_p_ =0.0019). If anything, the undershoot became larger with increasing levels of target force in young compared to old participants (Fig.3C). That is, this group of older participants performed at least as good if not better than their young counterparts in the slider condition.

The slider condition is linked to integrity of somatosensory function. This sensory function was also investigated by means of the position-matching task but this task (see methods) did not reveal any differences in somatosensory function between the young and older participants (supplementary note 3).

### Older participants exert higher forces in direct conditions than younger participants

The results from the slider condition and from the position-matching task suggest that our two age groups had similar somatosensory abilities. We then looked at differences in the direct conditions where sensory attenuation is supposed to happen.

In the mirror condition (Fig.3F), force errors in the young group were consistently more negative than those in the older group of participants. Mean force error of older participants was even positive for 4 and 6 N, indicating that the older participants produced more force than they should. Therefore, both young and older adults performed differently in the mirror condition with the magnitude of force errors being higher in the older adults (Analysis 1, main effect age F(1,68)=4.53, p=0.037, *η*^2^_p_ =0.49). With increasing levels of forces, the forces errors became more negative in the young participants and transitioned from positive to more negative in older ones (Analysis 1, main effect level of force, F(2, 136)=138.5, p<0.0001, *η*^2^_p_ =0.499). We could not find any evidence for a different scaling of force error with target force between the two age groups (Analysis 1, age x level of force, F(2, 136)= 2.47, p=0.088, *η*^2^_p_ =0.0089).

In the parallel condition (Fig 3I), the force errors of young group were positive for 4 N but negative 6 and 8N target forces. The mean force errors in older group was consistently positive for all levels of forces and was therefore higher (i.e. more positive) than the force errors of younger participants, indicating once again that the overshoot was larger in old compared to young participants (Analysis 1, main effect age F(1,68)=10.63, p=0.0017, *η*^2^_p_ =0.79). The force errors were again scaled with target force, becoming more negative with increasing levels of forces, the force errors became more negative in similar ways for both age groups (Analysis 1, main effect level of force, F(2, 136)=27, p<0.0001, *η*^2^_p_ =0.19). In this condition too, we could not find any evidence for a different scaling of force error with target force between the two age groups (Analysis 1, age x level of force, F(2, 136)= 1.37, p=0.26, *η*^2^_p_ =0.0099)

In each age group separately, the participants produced different amount of forces in the different conditions, with force errors being more positive in parallel, followed by mirror and by slider (Analysis 2, main effect condition: young: F(2, 68)=3.82, p=0.027, *η*^2^_p_ =0.31; old: F(2, 68)=16.75, p<0.0001, *η*^2^_p_ =0.69). This difference across conditions suggests that both groups exhibited some level of sensory attenuation. Similarly, the force errors became more negative with increasing level of target force for both groups (Analysis 2, main effect level of force: young: F(2, 68)=105.6, p<0.0001, *η*^2^_p_ =0.67; old: F(2, 68)=63.80, p<0.0001, *η*^2^_p_ =0.29). The scaling of the force errors with target force appeared to vary slightly across condition in both age groups (Analysis 2, level x condition: young: F(4,136)=2.64, p=0.036, *η*^2^_p_ =0.019; old: F(4,136)=1.95, p=0.1, *η*^2^_p_ =0.01).

We then directly compared the force errors between the two age groups directly. In addition to the influence of condition (Analysis 3, main effect condition F (2,136)=19.67, p<0.001, *η*^2^_p_ =0.345), level of force (main effect level of force, F (2,136)=150.7, p<0.001, *η*^2^_p_ =0.261) and interaction (Analysis 3, level x condition, F (4,272)=3.02, p=0.018, *η*^2^_p_ =0.006) that we already highlighted above for both groups separately, we found that the difference in force errors between the slider condition and the two direct conditions (mirror and parallel) was larger for old than for young participants (Analysis 3, age x condition, F (2,136)=4.17, p=0.017 *η*^2^_p_ =0.0733). In other words, while participants from both groups undershot the target force in the slider condition, older participants exhibit an overshoot in the mirror and parallel conditions for most target force levels (positive force errors) while younger participants kept undershooting the target force (negative force errors).

### Higher overcompensation in parallel than in the mirror condition

To quantify the amount of sensory attenuation more accurately, we decided to take into account inter-subject difference in somatosensation (measured by the performance in the slider condition). To do so, we computed the normalized overcompensation by subtracting force errors measured in the slider condition from the force-errors observed in the direct conditions (Fig.4).

**Fig 4:**
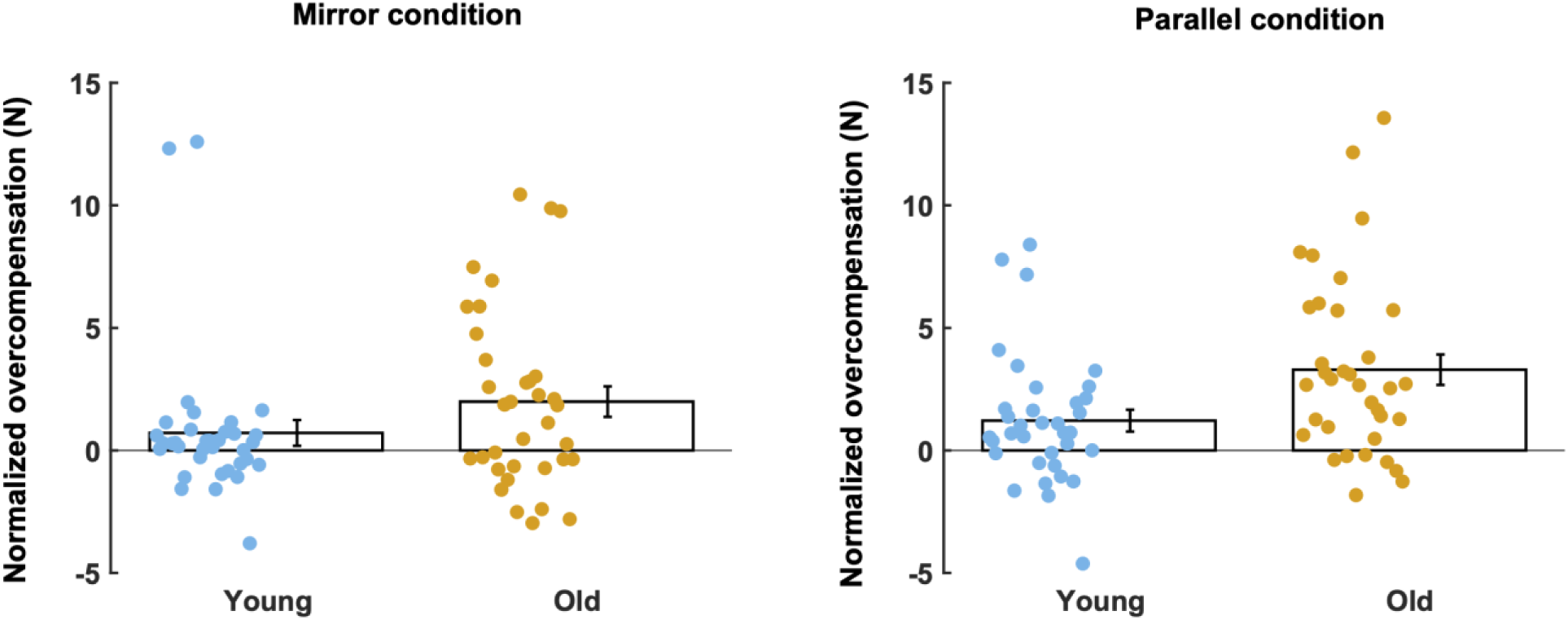
Normalized Overcompensation (experiment 1). The average normalized overcompensation is presented for each age group. (N=35/group) and each condition (panel A: mirror condition; panel B: parallel condition. Each dot is the average normalized overcompensation for each individual (collapsed across force levels). The black rectangle represents the average across all participants for each group separately. The error bar represents the standard error of the mean.

The mean normalized overcompensation was higher than zero in each direct condition for older adults but only in the parallel condition for younger adults (Analysis 4, young mirror: t(34)=1.359, p=0.1831 CI= [-0.354 1.786]; young parallel: t(34)=2.7278, p=0.01, C.I= [0.3107 2.1263]; older mirror: t(34)=3.2181, p=0.0028 CI= [0.73 3.325]; older parallel: t(34)=5.286, p<0.001 CI= [2.02 4.55]).

Older adults exhibited more sensory attenuation than young adults as showed by their higher mean normalized overcompensation (Analysis 4, main effect age F(1,68)=5.168, p=0.0262, *η*^2^_p_ =0.71). In addition, normalized overcompensation allows us to compare the amount of sensory attenuation between both direct conditions. The overcompensation was higher in the parallel than in the mirror condition (Analysis 4, main effect condition F(1,68)=10.02, p=0.0023, *η*^2^_p_ =0.2). However, we did not find any evidence that this difference between conditions was different for the two age groups (Analysis 4, age x condition F(1,68)=1.96, p=0.166, *η*^2^_p_ =0.04)

### Replication of higher sensory attenuation with age in the mirror condition

Explaining the task to the participants in experiment 1 was much harder than anticipated. We were therefore worried that some of the effects could be driven by the fact that the older participants did not understand the instructions correctly. Therefore, we redesigned the task training (see methods) and performed a replication of our mirror and slider conditions. In contrast to Experiment 1, we did not find any evidence that the young and older adults performed differently in the slider condition (Analysis 5, main effect of age: F(1, 59)=0.0109, p=0.9173, *η*^2^_p_ =0.0004). The force errors varied across the three levels of forces as they become more negative with increasing levels of target forces (Analysis 5, F(2,118)=99.91, p<0.0001, *η*^2^_p_ =0.99). For this group of participants, the results of the position-matching task (see methods) did not reveal any differences between the two age groups in the somatosensory abilities (supplementary note 3).

In the mirror condition, young and older participants exhibited different pattern of force error (Fig.5b). Older participants were mostly overshooting the target force (positive force error) while the younger participants undershot it (Analysis 5, F(1,59)=4.096, p=0.0475, *η*^2^_p_ =0.492). In addition, young and older adults showed difference in their force errors across the three levels of forces. Here again, a scaling effect difference was detected (Analysis 5, F(2,118)=91.4344, p<0.0001, *η*^2^_p_ =0.5).

**Fig 5:**
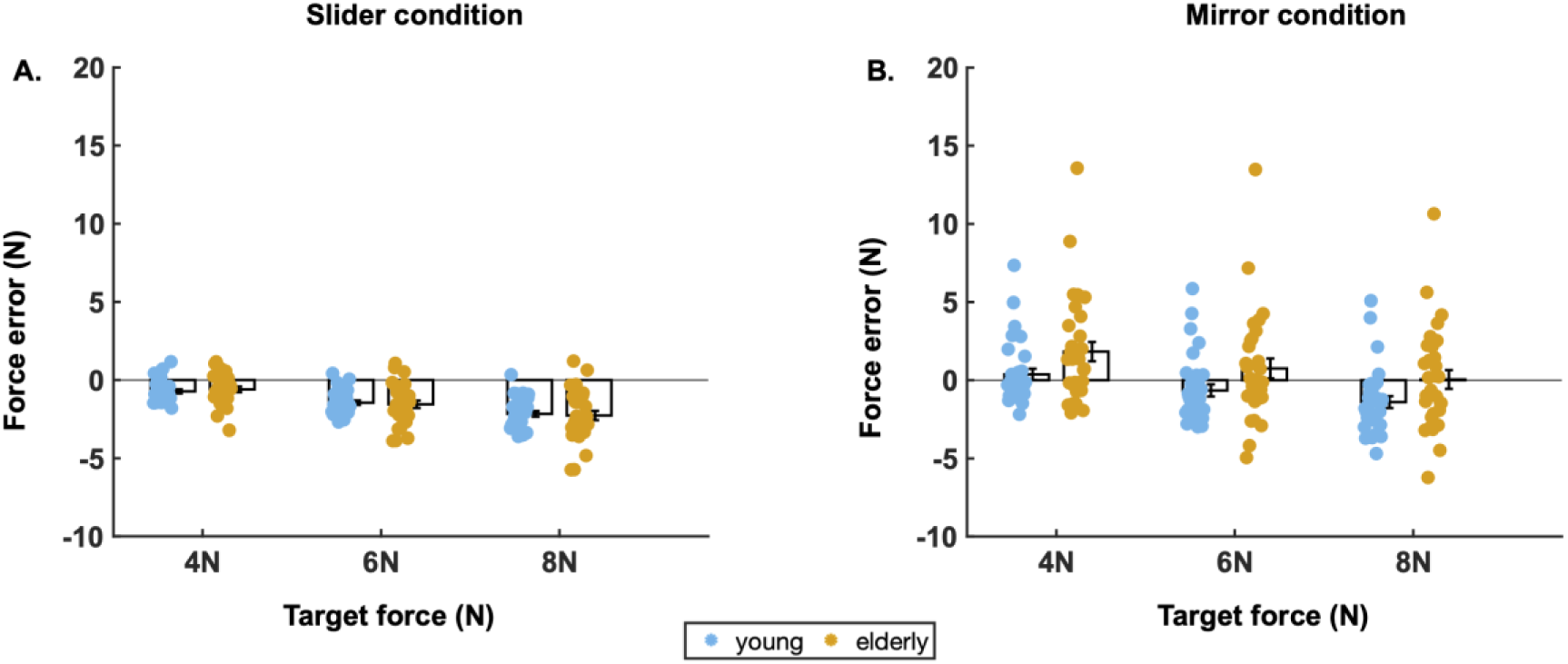
Comparison of force error between age groups (experiment 2). The average force errors (N=31 young, N=30 old) for both age groups and each force level for the slider condition (panel A) and mirror condition (panel B). Black rectangle and error bars represent the mean and standard error respectively. Each dot is the average across all trials for each force level and for each participant separately.

Young adults exhibited less undershoot in the mirror than in the slider condition (Analysis 6, main effect condition: F(1,30)=6.15, p=0.0189, *η*^2^_p_ =0.3096). The older participants even exhibit an overshoot in the mirror condition while they also undershot the target force in the slider condition (Analysis 6, main effect condition: 17.44, p<0.0001, *η*^2^_p_ =0.7335). For both age groups, force errors became more negative with increasing levels of forces (Analysis 6, main effect level of force: young: F(2,60)=90.88, p<0.0001, *η*^2^_p_ =0.68; old: main effect level of force F(2,58)= 60.7, p<0.0001, *η*^2^_p_ =0.2661). However, the difference in force errors across the two conditions (mirror vs. slider) was larger in older participants compared to their younger counterparts (Analysis 7, age x condition, F(1,59)=4.9002, p=0.03, *η*^2^_p_ =0.097).

As a result, the normalized overcompensation was positive for both young and old participants (Analysis 8, young: t(30)=2.48, p=0.0189, CI=[0.15, 1.61]; old: t(29)=4.17, p<0.001, CI=[1.2 3.5]) (Fig.6). Furthermore, this normalized overcompensation was larger for older compared to younger participants (Analysis 8, main effect age, F(1, 59)=4.9, p=0.0307, *η*^2^_p_ =0.9764). This confirms the results from our first experiment that sensory attenuation was higher in older than younger adults.

**Fig 6.**
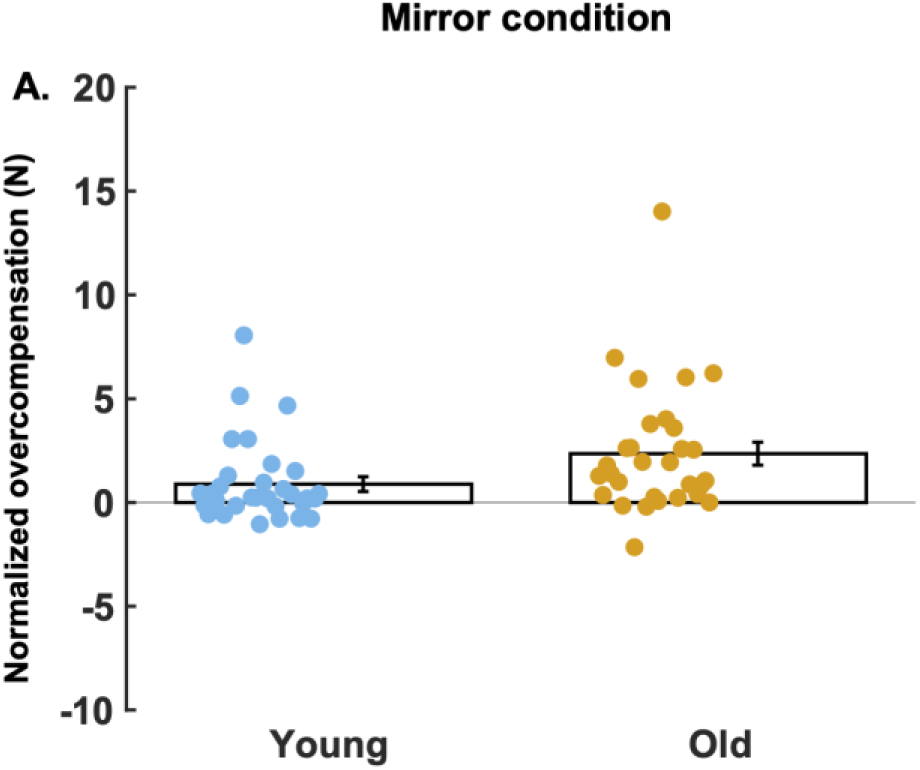
Normalized Overcompensation (experiment 2). The average normalized overcompensation is presented for each age group. (N=31 young, N=30 old) in the mirror condition. Each dot represents the average normalized overcompensation for each individual (collapsed across force levels). The black rectangle represents the average across all participants for each group separately. The error bar represents the standard error of the mean.

## Discussion

In our study, we found that both young and old participants exerted higher forces in the direct conditions (self-produced forces) than in the slider condition but this overcompensation was even higher in older participants. We did not find any evidence that force reproduction in the slider condition or accuracy in a position-matching task (which were our proxies for somatosensory function) were affected by age. While an increase in sensory attenuation with age (Klever et al. 2019; Wolpe et al. 2016) had been observed for the fingers, we confirm that this phenomenon generalized to another effector: the arm.

### Higher reliance on internal forward models with aging

By normalizing our data with respect to the slider condition, we were able to remove the influence of the somatosensory component and to isolate sensory attenuation. Our findings show that older participants had a higher sensory attenuation than younger adults did. An advantage of increased attenuation could indicate a preservation of a sense of agency or the sense that one controls one’s own actions and their consequences (Wolpe et al. 2016). Reduced sensory attenuation has been linked to impaired awareness of action and disorders such as schizophrenia and psychogenic movement disorders (Shergill et al. 2003; Wolpe et al. 2016).

The observed increase in sensory attenuation from this study together with their supposed age-related decline in sensory function (Dunn et al., 2015, Goble et al. 2009) suggests that elderly adults might rely more on the internal models (Wolpe et al. 2016). When a perceived force is self-produced, the sensation is a combination of sensory information with the predicted sensory consequences of the force generation (coming from the internal model). These signals are combined via Bayesian integration in function of their reliability (Körding et al. 2004; Orban de Xivry et al. 2013). Given that proprioceptive input becomes less reliable with increasing age, the weight of the internal model (which is shown not to be impaired by aging) becomes larger (Wolpe et al. 2016). In other words, there is a higher weighting on the internal model during the parallel processes involving both the internal model and sensory system when making a prediction (Bubic et al. 2010). Many studies have shown aging changes weighting on sensorimotor predictions during movements (Klever et al. 2019; Moran et al. 2014). Studies also have shown internal forward model function does not change with aging (Heuer et al. 2011; Vandevoorde and Orban de Xivry 2019).

While there is a shift towards higher sensory attenuation with aging, a shift in the opposite direction is observed in cerebellar patients or in people with schizophrenia (Knolle et al. 2013; Shergill et al. 2005). This shows that sensory attenuation is the outcome of an adaptable combination between sensory and predictive signals in function of their reliability such as been observed in different contexts (Bogadhi et al. 2013; Deravet et al. 2018; Ernst and Banks 2002; Orban de Xivry et al. 2013). The increase in sensory attenuation with aging shows that the reliability of the sensory signal decreases faster with aging than the reliability of the internal model signal (Wolpe et al. 2016).

### We did not observe an age-related decrement in somatosensory function

It remains to be understood why older participants assign a higher weight to their internal predictions while we did not find any impairment in sensory function. Indeed, we could not find any age-related differences in either the slider condition or the position-matching task in our samples.

Previous studies show that sensory function and proprioceptive acuity decrease with aging (Dunn et al. 2015; Goble et al. 2009; Ranganathan et al. 2001). In the slider condition, participants only had to indicate the perceived force on a slider; there were no self-produced forces. This condition provides us with a proxy for somatosensory function. Our results from both experiments show that both young and elderly adults undershot the target forces. In experiment 1, this undershoot was larger for the young participants. Hence, elderly participants were more accurate, i.e. their produced forces were closer to target forces. However, in experiment 2, we did not observe such age-related difference. In contrast, young adults from the study of Wolpe et al. (2016) were on average less accurate than the older adults but, in contrast to our experiment, they overshot the target forces.

Both our experiments and that from Wolpe et al. used different level of target forces. In Wolpe et al., the older participants scaled their produced forces with target force less accurately than the younger participants did. In our study, we found that both age groups exhibited a larger undershoot with increasing levels of target force. However, we failed to find any evidence for an effect of age on the scaling of the produced force with the target force in the slider condition in both experiments. Walsh et al. (2011) report a similar finding in their finger force matching experiment that subjects overestimated smaller target forces than larger ones. Their matched forces were 2-3 N higher than the smaller target forces.

Overall, across the 130 participants, there were no differences in somatosensory processing between the two age groups as opposed to what was shown by previous studies (Dunn et al. 2015; Goble et al. 2009; Morrison and Newell 2012; Ranganathan et al. 2001). This can maybe attributed to the fact that our participants were probably more active than general population of the age groups between 55-75 years. In addition, our mean age in the elderly (mean=64 years) was lower than the mean age of eldery in previous studies (mean= 71 years, Goble et al, Doumas et al 2008).

Yet, while our older participants did not exhibit any impairment in somatosensation, they did exhibit increased sensory attenuation. This begs the question whether the observed increase in sensory attenuation is due to poor sensory function as suggested by Wolpe et al. 2016. It rather seems to violate the idea that the increase in sensory attenuation is due to a shift in reliability-based balance between predictive and sensory signals. One possibility is that the average performance is similar but that the confidence in the sensory estimates is degraded with aging. Unfortunately, none of our somatosensory tasks have enough repetitions (maximum 10) to measure standard deviation in a reliable way.

### Sensory attenuation is higher in parallel condition than in the mirror

We used two different direct conditions where the force exerted by the right arm was directly felt on the left arm. In the mirror condition, the biceps of the right arm and the triceps of the left arm were simultaneously activated (non-homologous muscles). In contrast, in the parallel condition, both arms’ biceps muscles (homologous muscles) were simultaneously activated. Our results show that sensory attenuation is higher when homologous muscles are activated.

Humans naturally couple limb movements and it is usually easier to move limbs in the same direction and contract homologous muscles (Huang and Ferris 2009; Meesen et al. 2006, Baldiserra et al., 1982). In addition, humans identify and perceive forces applied by the hand in terms of the motor activity required to resist or produce the force or a “sense of effort” rather than in terms of a perceived force magnitude (Toffin et al. 2003). In other words, force perception is controlled by the ease of resisting it rather than the actual force magnitude (Van Beek et al 2013). Given the more natural connections between homologous muscles, we postulate that the sense of effort required to contract homologous muscles was felt as lower in the parallel condition, which could explain the larger overshoot in this condition if one tries to match the effort perceived in the force perception phase.

In our study, there activation of the homologous muscles was coupled to a change in direction of the force in the left arm. While we are confident that it does not largely affect the amount of overcompensation, this could even become larger if the force direction was not changed. A future study could reproduce this result in the absence of change of force direction.

### Limitations

While we have shown the effect of sensory attenuation on the arms, further investigation is required on the age-related differences between the mirror and parallel conditions. In the parallel condition, there was both an activation of homologous muscles and change in direction of movement. We believe that this change in direction did not play a big role in the result of the present study. Yet, a future study could correct this mistake and involve pushing the right arm in the rightward direction and the transmission in the rightward direction. Further studies on somatosensation on the arm are required to explore the age-related differences or similarities deeper.

In addition, the understanding of the task instructions by the participants could have had an influence on the performance. We overcame this limitation in experiment 2 with clearer task instructions. Nevertheless, participants in both experiments reported that the task was difficult to follow, and future studies must be done with careful consideration to the development of detailed task instructions.

Moreover, the participants in our study was limited in number, and we divided them into two arbitrary age groups (N=35 for each of them in experiment 1 and N=30/group in experiment 2). Future studies are warranted to include a larger participant pool from a larger age range, such that it will also be possible to conduct correlation analyses between age and behavioral indices.

### Conclusion

Our force-matching paradigm sheds a new light on the effect of aging on sensory attenuation and somatosensation. First, we confirmed that sensory attenuation can be observed in the arms, similar to what has been found for the fingers (Logan et al. 2019; Shergill et al. 2003; Walsh et al. 2011; Wolpe et al. 2016) and that this is a phenomenon that goes beyond the fingers. Second, we replicated the finding that sensory attenuation is larger in adults over 55 years. The Bayesian perspective adopted by Wolpe et al. would let us interpret these results as indicative of a shift in the balance between sensory and predictive signals, which is in line with the hypothesis that internal model function is unaffected by aging (Heuer et al. 2011; Vandevoorde and Orban de Xivry 2019). Yet, in our sample, we did not detect any difference in somatosensory function between our two age groups. This leads us to question the fact that the increased sensory attenuation with age is due to a deficit in proprioception.

## Supplementary information

All supplementary information, raw data, processed data and scripts are available <u>here</u>.

## Acknowledgements

This work was supported by an internal grant of the KU Leuven (STG/14/054) and by the FWO (Hercules I005018N). We thank Remie De Crock, Olivier De Weer and Astrid Mulier for helping us with the data collection

